# UFMylation of MRE11 is essential for maintenance of telomere length and hematopoietic stem cell survival

**DOI:** 10.1101/846477

**Authors:** Lara Lee, Ana Belen Perez Oliva, Dmitri Churikov, Elena Martinez-Balsalobre, Joshua Peter, Dalicya Rahmouni, Gilles Audoly, Violette Azzoni, Stephane Audebert, Luc Camoin, Victoriano Mulero, Maria L. Cayuela, Vincent Geli, Yogesh Kulathu, Christophe Lachaud

**Author notes:** **Correspondance** Christophe Lachaud, Aix-Marseille Univ, INSERM, CNRS, Institut Paoli-Calmettes, CRCM, Marseille, France. Equipe labellisée Ligue (Dmitri Churikov, Vincent Geli). Equal contribution (Lara Lee, Ana Belen Perez Oliva).

## Abstract

Genetic studies using knockout mouse models provide strong evidence for the essential role of the ubiquitin-like protein UFM1 for hematopoiesis, especially erythroid development, yet its biological roles in this process are largely unknown. Here we have identified a UFL1-dependent UFMylation of the MRE11 nuclease on the K281 and K282 residues. We show that Hela cells lacking the specific UFM1 E3 ligase display severe telomere shortening. We further demonstrate either by deleting UFM1 or by mutating MRE11 UFMylation sites that preventing MRE11 UFMylation impacts its interaction with the telomere protein TRF2. However, the MRE11 function in double-strand-break repair remains intact. We validate these results *in vivo* by showing that Zebrafish knockouts for the genes *ufl1* and *ufm1* have shorter telomeres in hematopoietic cells. Here we present UFMylation has a new mechanisms of regulation for telomere length maintenance with a role in hematopoiesis.

**Key points:** Modification of MRE11 by UFM1 regulates telomere maintenance and cell death in HSCs

**Scientific category:** UFMylation, telomere maintenance, hematopoietic stem cell survival.

## Introduction

The ubiquitin family proteins are composed of Ubiquitin (Ub), and the other ubiquitin-like (Ubl) proteins, all of which share the common β-grasp fold. They are posttranslational modifiers that play pivotal roles in a diverse range of cellular processes. These include cell cycle progression, DNA damage response, protein translation and stability, signal transduction, intracellular trafficking, and antiviral response {Hochstrasser:2009hr}.

Ub-fold modifier 1 (UFM1) is one of the most recently identified Ubls. It is evolutionarily conserved, and its orthologs can be found in Metazoa and plants, but not in yeast (Komatsu *et al*, 2004). The process of protein modification by UFM1 involves an enzymatic cascade similar to those described for the other ubiquitin family members. Briefly, after cleavage of Pro-UFM1 by the peptidase UFSP2, UFM1 is activated by the Ufm1-specific E1 enzyme UBA5, conjugated by the E2 enzyme UFC1 and ligated by the E3 ligase UFL1. UFM1 is cleaved from its targets by the UFM1-specific protease, UFSP2 (reviewed in (Daniel & Liebau, 2014)).

Genetic studies using knockout mouse models and the identification of patients with mutations in the UFM1 pathway provided strong evidence for the indispensable role of this posttranslational modification in animal development and homeostasis. Indeed, it has been shown that UFM1 is crucial for hematopoiesis (Tatsumi *et al*, 2011; Zhang *et al*, 2015; Cai *et al*, 2015), liver development (Yang *et al*, 2019), brain development (Nahorski *et al*, 2018), heart failure protection (Li *et al*, 2018), maintenance of intestinal homeostasis, and protection from inflammatory diseases (Cai *et al*, 2019)..

The nuclear receptor co-activator ASC1 (Activating signal co-integrator 1) has been one of the first UFMylated protein identified. UFMylated ASC1 enhances the recruitment of transcription co-factors to promoters of estrogen receptor α (ERα) target genes and upregulates their expression (Yoo *et al*, 2014). More recently, the ribosomal subunit RPL26 has been suggested to be the principal target of UFMylation. Its UFMylation is suggested to mediate ER homeostasis by regulating protein biogenesis of secretory proteins (Walczak *et al*, 2019). Additionally, upregulation of the protein levels of p53 and p53 targets, as well as increased level of H2AX phosphorylation have been reported in Ufl1-deficient bone marrow cells (Zhang *et al*, 2015). This suggests that the UFM1 pathway could contribute to genome stability, although the exact mechanism whereby Ufl1 exerts this function remains unknown. Very recently, it was reported that UFM1 promotes ATM activation and monoufmylates histone H4 in response to double strand breaks suggesting that H4 UFMylation contributes to the amplification of ATM activation (Qin *et al*, 2019).

In this study, we systematically analyzed the functions of UFMylation in genome stability by focusing on the E3 ligase UFL1. Our work reveals a hitherto unanticipated role for UFMylation in maintaining telomere length. We further define the underlying mechanism by identifying that modification of MRE11 with UFM1 is crucial for telomere maintenance in HeLa cell lines. Consistent with this result, loss of UFMylation resulted in marked shortening of telomeres during hematopoiesis in animal models. These findings provide a likely explanation for the observed anemia in mice lacking components of the UFM1 pathway.

## Results

### UFL1 localizes to the nucleus and chromatin

UFM1 was initially detected within the nucleus (Komatsu *et al*, 2004) however, UFL1 has been described as a protein localized to the endoplasmic reticulum (ER) via its interaction with the ER resident protein UFBP1 (Zhu *et al*, 2019). We thus investigated the localization of both UFL1 and UFM1 using cell protein fractionation assay in HeLa cells. In this experiment we detected UFL1 in the cytoplasmic (CET), nuclear (NEB) and chromatin (NEB+) fractions. Whereas free (unconjugated) UFM1 appeared to be predominantly cytoplasmic, UFMylated proteins were detected in the nucleus and chromatin (Fig.S1A). We also analyzed the localization of UFM1 and UFL1 by expressing GFP tagged forms of these proteins. We detected both proteins in the cytoplasm, nucleus, and the chromatin under unstressed conditions (Fig. S1B). Since most of the proteins involved in DNA repair are recruited to chromatin following DNA damage, we tested whether UFL1 localizes to chromatin after ionizing irradiation (IR) using subcellular protein fractionation. Indeed, we observe a strong enrichment of UFL1 after IR at the chromatin, suggesting that UFL1 may be involved in DNA repair (Fig. S1C).

### Loss of UFL1 results in defective telomere maintenance

To get insights into the function of UFL1 in DNA repair we next used CRISPR/CAS9-mediated genome editing to disrupt the *UFL1* gene in HeLa cells (Fig. S2A,C). Analysis of genomic DNA from clonal populations of HeLa cells obtained using a gRNA targeting exon 2 revealed a deletion and frame shift in both alleles (Fig. S2D). Consistent with published results (Zhang *et al*, 2015), we noticed that UFL1 KO cells have a reduced growth rate (Fig. S2E). This latter result prompted us to investigate whether UFL1 KO cells were defective in DNA repair and exhibited chromosome abnormalities. Interestingly, chromosome spreads experiments revealed that UFL1 KO cells displayed marked increased levels of spontaneous chromosomal fusions compared to WT HeLa cells (Fig. 1A left panel, B). Because DNA-repair activities at deprotected telomeres can generate chromosome end-to-end fusions (Maciejowski & de Lange, 2017), we assessed the effect of UFL1 inactivation on telomere integrity by using fluorescence in situ hybridization (FISH) on metaphase chromosome spreads. This approach revealed four times more chromatids ends without detectable telomeric signal in UFL1 KO cells compared to WT HeLa cells (Fig. 1A right panel, B) indicating that loss of UFL1 increases the frequency of telomere losses.

**Figure 1:**
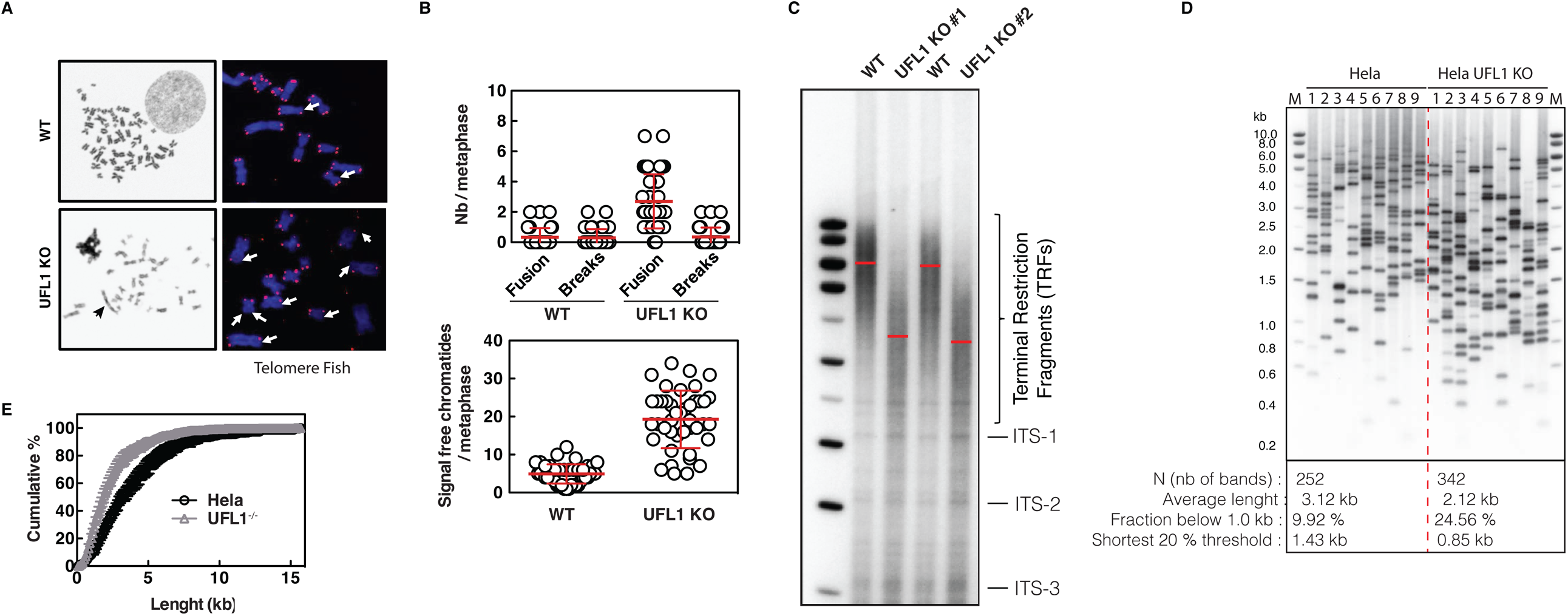
UFL1 KO cells have defect in telomere length. (A) Representative images of telomere FISH in Hela and Hela UFL1KO cells. Blue, DAPI-stained chromosomes. Red dots, telomeres; White arrows, telomere loss. (B) Histogram shows quantification of chromosomes breaks and fusion (top histogram) and signal free chromatids ends per metaphase (Bottom histogram). The quantification have been done on 40 metaphase from 2 different experiments. (C) Southern blot analysis showing TRF distributions from indicated cells. Red line indicates the size of the TRF bands. (D) Results of TeSLA using DNA as indicated. Nine TeSLA PCRs were performed for each DNA sample. (E) Quantification of D.

To verify whether UFL1 KO results in global shortening of telomeres, we performed Southern blot analysis of terminal restriction fragments using two independently derived UFL1 KO clones. Both clones showed drastic decrease of the mean telomere length compared to the WT controls (Fig. 1C). Taken together these results suggest an important role for UFL1 in telomere maintenance. To further characterize the changes in telomere length in more detail, we used the Telomere Shortest Length Assay (TeSLA) that reveals the length of individual telomeres from all chromosome ends (Lai *et al*, 2017). These analyses reveal that the fraction of very short telomeres (< 1 kb) is increased more than two-fold in UFL1 KO compared to WT HeLa cells (Figs 1D and S3). The TeSLA analysis revealed up to 40% reduction in the average length of amplified telomeres in UFL1 KO clones further highlighting the importance of UFL1 in telomere length regulation.

Taken together, the results of Figure 1 show that loss of UFL1, the only known E3 ligase for UFM1 conjugation to substrates, results in telomere length shortening and telomere loss unveiling a role for UFMylation in telomere maintenance.

### MRE11 interacts with the UFM1 pathway members

In order to understand how UFL1 regulates telomere length we sought to identify new interactors of UFL1. The identification of enzyme partners has long been challenged by the transient character of the interaction that does not always allow purifying the complex by conventional protein co-immunoprecipitation methods. To overcome this technical issue, we used proximity labelling using the BioID method that biotinylates proteins in close vicinity (Kwon & Beckett, 2000). Biotinylated proteins can then be purified using streptavidin affinity capture and identified by mass spectrometry. To identify substrates and interactors of the UFM1 E3 ligase UFL1, we fused BirA (R118G) biotin ligase domain to the N-terminus of UFL1 (Kwon & Beckett, 2000). The BirA-UFL1 fusion protein was inducibly expressed in Hela FRT-TREx cells. Cells expressing BirA fused to GFP were used as a negative control. Doxycycline concentration and treatment time was titrated to minimize overexpression artefacts (Fig. 2A). We verified that the fusion of BirA on UFL1 did not alter the interaction of UFL1 with its known partner UFSP2. As expected, UFSP2 is only detected in the biotin pull-down from BirA-UFL1 expressing cells (Fig. 2B). After validation of the cell lines, we performed mass spectrometry to identify the biotinylated proteins. Proteins unique to the BirA-UFL1 pull-down, and not detected in the BirA-GFP pull-downs were considered for further analysis (Fig. 2C). As expected, mass spectrometry identified the known partners of UFL1 such as CDK5RAP3, UFBP1 and UFSP2. Interestingly, using this approach we identified MRE11 as a protein biotinylated by BirA-UFL1 (Fig. 2C). To test if MRE11 is also an interactor of UFM1 and UFC1, we tested biotinylation in cells expressing BirA-UFM1 or BirA-UFC1, which revealed MRE11 to also be an interactor of UFM1 and UFC1 (Fig.S4). We further confirmed the presence of MRE11 in biotin pulldown from BirA-UFL1 (Fig. 2D) and BirA-UFM1 expressing cells (Fig. 2E) by immunoblotting. MRE11 together with RAD50 and Nijmegen breakage syndrome 1 (NBS1; also known as nibrin) forms the multifunctional protein complex MRN. Besides its role in the maintenance of genome stability (Stracker & Petrini, 2011), MRN has been also reported to promote C-NHEJ at dysfunctional telomeres through the activation of ATM (Attwooll *et al*, 2009; Deng *et al*, 2009) (Dimitrova & de Lange, 2009). The MRN complex has also been proposed to regulate telomere length (Wu *et al*, 2007) (Cai *et al*, 2015) suggesting a possible link to telomere maintenance by UFMylation.

**Figure 2:**
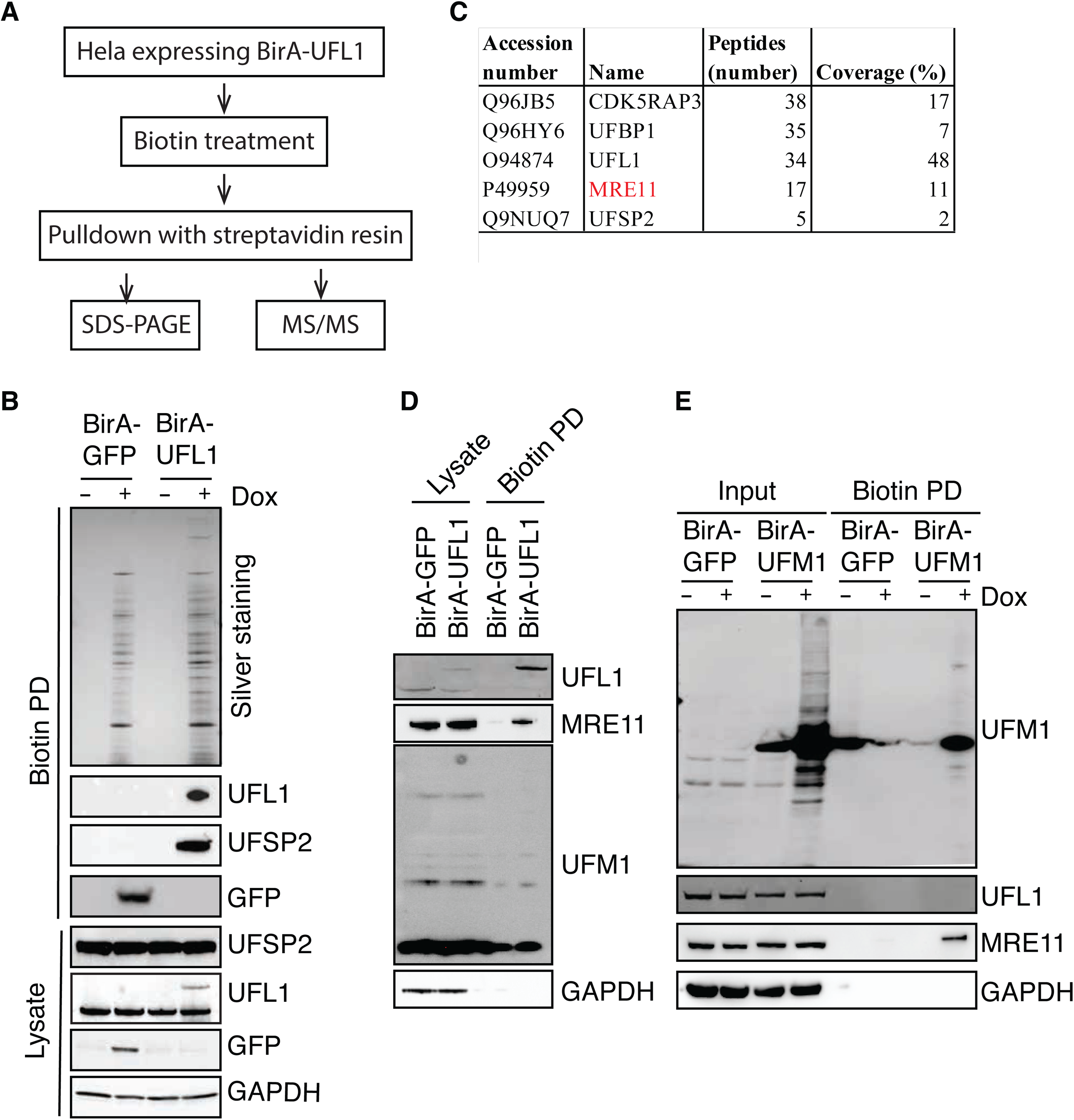
UFL1 interacts with MRE11. (A) Strategy for the identification of UFL1 binding proteins. (B) Biotinylated proteins are purified from cells expressing BirA-GFP or BirA-UFL1. The samples were then subjected to immunoblot with indicated antibodies (C) List of the candidate target proteins identified by mass spectrometry. (D) Biotinylated proteins are purified from cells expressing BirA-GFP or BirA-UFL1. The samples were then subjected to immunoblot with indicated antibodies. (E) Biotinylated proteins are purified from cells expressing BirA-GFP or BirA-UFM1. The samples were then subjected to immunoblot with indicated antibodies.

### MRE11 is a substrate of UFL1

We next investigated whether MRE11 is a substrate of UFL1. We therefore reconstituted the UFMylation enzyme cascade *in vitro* using purified UFM1 and the enzymes UBA5, UFC1 and UFL1. When recombinant MRE11 was added into reactions, a band shift of ∼10kDa was detected corresponding to the addition of a single UFM1 was observed (Fig; 3A). Importantly, this modification is dependent on the presence of UFL1, suggesting that MRE11 is a direct substrate of UFL1 (Fig 3A). To test if MRE11 is also UFMylated in cells, we analyzed cell extracts from Hela cells with an Mre11 antibody. In accordance with the *in vitro* reconstitution results, a similarly shifted band of MRE11 was detected in extracts prepared from WT cells but not UFL1 deficient cells (Fig. 3B). While UFL1 is known to only mediate the attachment of UFM1 to proteins, the observed loss of modified MRE11 in UFL1 KO cells could be an indirect effect and MRE11 could be modified by a different PTM. To confirm that the higher molecular weight band of MRE11 detected is indeed UFMylated MRE11, we incubated the protein extracts from WT cells with recombinant UFSP2. Indeed, incubation with UFSP2 results in the disappearance of the slower migrating band confirming that the observed band is UFMylated MRE11 (Fig. 3B). Taken together these results demonstrate that UFL1 UFMylates MRE11.

**Figure 3:**
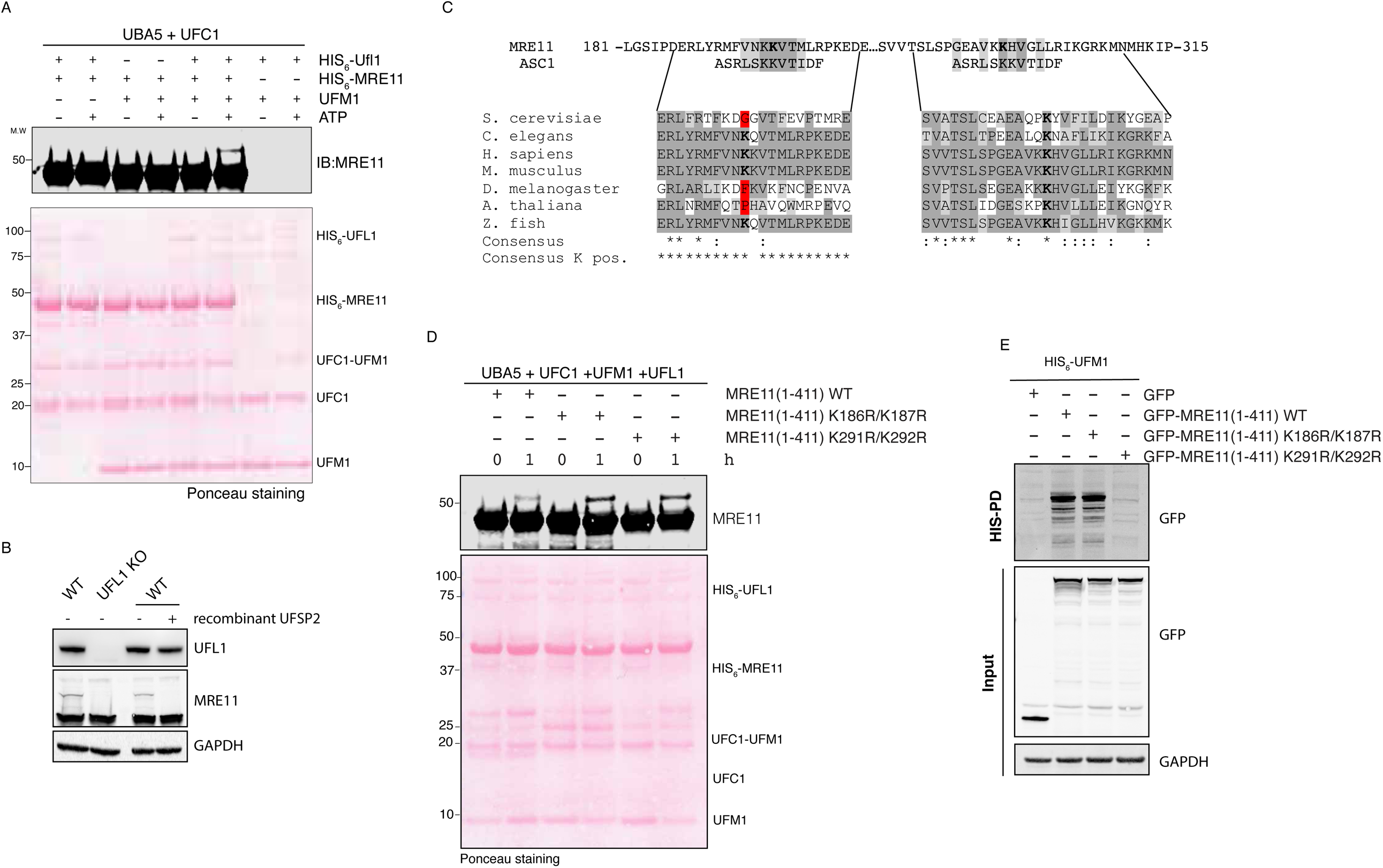
MRE11 is Ufmylated on K296-297. (A) Recombinant His6-MRE11 were incubated with Uba5, Ufc1, Ufl1 and Ufm1 in the presence of ATP and MgCl2 for 1 hr at 37°C. The reaction was stopped by addition of 3xSDS loading buffer and the reaction products were separated on a 4-12% NuPAGE SDS-PAGE gel. Top panel - Western blot analysis of ufmylation of MRE11 WT and mutants using MRE11 specific antibody. Bottom panel - Ponceau stained nitrocellulose blot as loading control. (B) 10^6^ cells are lysed in SDS and after treatment or not with recombinant UFSP2, samples were subjected to immunoblot with indicated antibodies. (C) Protein alignment of ASC1 ufmylated site with MRE11 sequence. (D) Same as A with MRE11 fragment mutated in indicated residues. (E) GFP tagged MRE11 WT and mutant were expressed in cells with 6HIS-UFM1 as indicated. Cell lysates were subjected to pull-down with NTA resins followed by immunoblot with indicated antibodies.

We next wanted to identify the lysine residues in MRE11 that get UFMylated. MRE11 contains 45 lysine residues, and we wondered if we could define a consensus UFMylation site based on homology to the identified UFMylation sites on ASC1 (Fig. 3C). Using this approach, we identified four lysines within two stretches of MRE11 (K196-K197 and K281-K282). To test if these lysines in MRE11 are the preferred UFMylation sites, we mutated each pair to arginine. Whereas, mutation of the candidate lysines to arginines did not affect the ufmylation of MRE11 fragment *in vitro* (Fig 3D), K281R, K282R mutant showed a strongly reduced UFMylation when expressed in HeLa cells (Fig. 3E). This result suggests that MRE11 is UFMylated on K281-K282 but possibly also on other lysine residues.

### UFL1 does not regulate MRE11 function in DSB repair and HR

MRE11 is part of the MRN complex, a sensor of DSBs that also controls the DNA damage response (DDR) by governing the activation of the central transducing kinase ataxia-telangiectasia mutated (ATM). In addition, the MRE11 complex regulates DSB repair, through the homology directed repair (HDR) (review in (Oh & Symington, 2018)). In order to investigate if UFL1 controls MRE11 functions in the DDR, we analyzed whether the interaction between MRE11 and UFM1 pathway members was dependent on DNA lesions. Mass spectrometry analysis of the proteins interacting with UFM1 (Fig. S4A,D,E), UFC1 (Fig. S4B,D,E) and UFL1 (Fig. S4C,D,E) showed that exposure of cells to irradiation did not increase the biotinylation of MRE11. This suggests that the interaction between MRE11 and the UFL1 pathway is not increased upon DNA damage. These results were further confirmed by immunoprecipitation of MRE11 from cells following DNA damage. As indicated in Fig 4A, the amount of UFL1 found associated with MRE11 did not change after induction of DSB by irradiation; and MRN complex formation was also not impacted in UFL1 KO cells (Fig. 4A). We then addressed whether UFL1 regulated the recruitment of the MRN complex to chromatin. Indeed, subcellular fractionation revealed that recruitment of the MRN complex to chromatin is increased in UFL1 KO cells (Fig. 4B). Increased levels of MRE11 at chromatin were confirmed by immunofluorescence on the chromatin fraction (Fig. 4C). One possible explanation for the recruitment of MRE11 to chromatin could be due to increased levels of DNA lesions in UFL1 KO cells. However, we found that γH2AX and pATR levels were similar in untreated WT and UFL1 KO cells (Fig. 4D, E), whereas phosphorylation of ATM was slightly enhanced in UFL1 KO cells (Fig. 4D). Finally, we investigated whether UFL1 could control DSB repair mediated by MRE11 by monitoring the activation of ATM and ATR as well as the phosphorylation of H2AX upon DNA damage. Compared to wildtype cells, phosphorylation of ATR, ATM (Fig. 4D) and H2AX (Fig. 4E) was slightly enhanced in UFL1 KO cells exposed to various DNA damaging agents compared to similarly treated WT cells. Additionally, the sensitivity of UFL1 KO cells to X-rays was not increased significantly compared to control cells (Fig. 4F). Since homologous directed repair (HDR) is also regulated by MRE11 and is necessary for the repair of lesions induced by X-rays we analyzed if defect in UFL1 impact HDR. Consistently with the absence of sensitivity of UFL1 KO cells to X-rays, GFP reporter assay indicates that HDR was not compromised in UFL1 KO cells (Fig. 4G). Taken together, these results demonstrate that UFL1 regulates neither DSB repair nor HDR mediated by MRE11.

**Figure 4:**
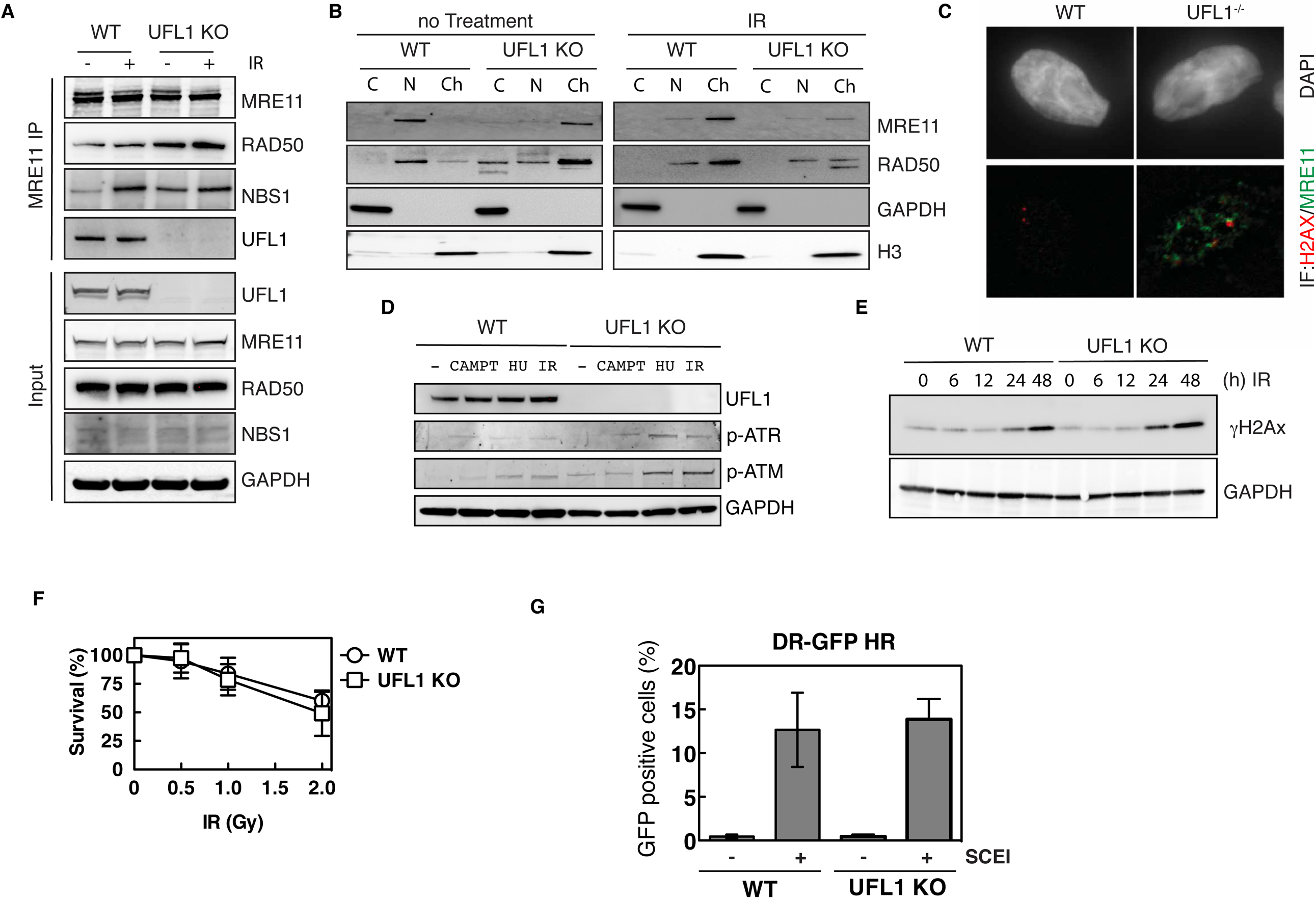
UFL1 does not regulate MRE11 function in DSB repair and HR. (A) Protein lysates from indicated cells treated or not with IR were subjected to MRE11 immunoprecipitation followed by immunoblot with indicated antibodies. (B) Immunoblot of MRE1, RAD50, H3 and GAPDH in the absence or presence of IR. C = cytoplasm fraction, N = Nuclear fraction and Ch= chromatin fraction. (C) MRE11 and ϒ-H2AX immunostaining of WT and UFL1 KO cells. (D) Immunoblot of UFL1 p-ATM p-ATR and GAPDH of protein extract from cells exposed to the indicated treatments. (E) Detection of ϒ-H2AX activation at various time post IR of WT and UFL1 KO cells. (F) UFL1 WT or KO cells exposed to various intensity of IR, were subject to clonogenic survival assays. For each cell population, viability of untreated cells is defined as 100%. (G) WT and UFL1 KO cells were co-transfected with HR reporter and pCBA-I-SCEI plasmids. The percentage of GFP positive cells represent cells active for HR.

### UFM1 regulates MRE11 TRF2 interaction

As the most striking phenotype of UFL1KO cells is telomere shortening, we addressed whether UFL1 could specifically regulate MRE11 function at telomere. The MRE11 complex localizes to mammalian telomeres via its interaction with the telomeric protein TRF2 (Zhu *et al*, 2000). We therefore analyzed if the interaction between MRE11 and TRF2 was modified in UFL1 KO cells. We first used a proximity ligation assay (PLA) to monitor proximity between MRE11 and TRF2. Interestingly, we observed a significant decrease in the number of foci detected by PLA in UFL1 KO cells compared to control cells (Fig. 5A, B). This suggests an UFL1 dependent interaction between MRE11 and TRF2. To confirm this result, we analyzed MRE11 immunoprecipitates from wildtype and UFL1 KO cells for association between MRE11 and TRF2. While similar amounts of RAD50 co-precipitated with MRE11 in UFL1 KO and control cells, the association of TRF2 was strongly reduced in the absence of UFL1 (Fig. 5C). The results of both PLA and co-IP indicate that the interaction between TRF2 and MRE11 is markedly reduced when UFL1 is inactivated and suggests that UFMylation of MRE11 is necessary for its interaction with TRF2. To confirm this hypothesis, we expressed GFP tagged versions of MRE11 mutants and tested their association with TRF2. Whereas TRF2 co-immunoprecipitated with GFP-MRE11 and GFP-MRE11 K196-7R mutant, the UFMylation defective GFP-MRE11 K282-3R mutant barely associated with TRF2. Importantly, this mutant retained the ability to bind RAD50 confirming that mutating these lysines does not disrupt MRE11 folding or other functions (Fig. 5D).

**Figure 5:**
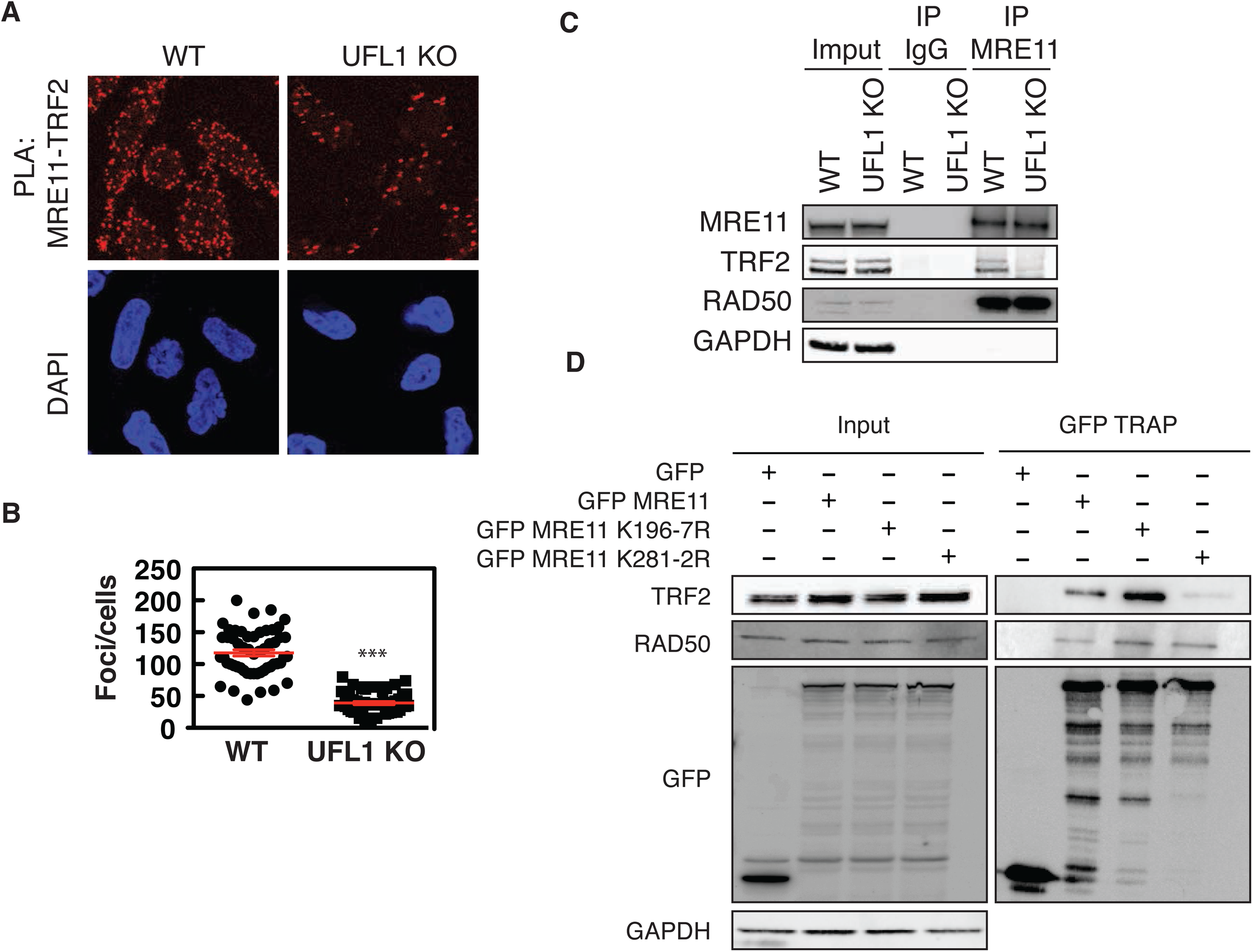
UFM1 regulates MRE11 TRF2 interaction. (A) In cells proximity ligation assay (PLA) with anti-TRF2 and anti-MRE11 antibodies in combination in WT and UFL1 KO cells. (B) Quantification of A. (C) Protein lysates from indicated cell were subjected to MRE11 immunoprecipitation. Immunoprecipitation with IgG was used as a negative control. After SDS PAGE, samples were analyzed with the indicated antibodies. (D) GFP tagged MRE11 WT and mutant were expressed in cells as indicated. Cell lysates were subjected to GFP trap followed by immunoblot with indicated antibodies.

In summary, these results show that interaction of MRE11 with TRF2 requires UFL1 and MRE11 UFMylation at lysine K282 and K283.

### UFM1 pathway is essential to prevent telomere instability and HSC death *in vivo*

As UFM1 has been shown to be crucial for hematopoiesis (Tatsumi *et al*, 2011; Zhang *et al*, 2015; Cai *et al*, 2015), we then studied the relevance of MRE11 UFMylation *in vivo* in a zebrafish model using the unique advantages of zebrafish for genetic analysis of hematopoiesis. As in mammals, zebrafish hematopoiesis occurs in two waves: the primitive wave generates a transient population of blood cells, while the second wave generates hematopoietic stem cells (HSCs) by around 30 hours post-fertilization (Robertson *et al*, 2016). Genetic inactivation of either *ufm1, ufl1* or *mre11a* with CRISPR-Cas9 (Fig. S5A-C) resulted in reduced erythrocyte numbers after 4 days post-fertilization (dpf) larvae (Fig. 6A, B) assayed in the zebrafish line *Tg(lcr:eGFP)* which expresses GFP in erythroid cells (Ganis *et al*, 2012). Notably, robust telomere shortening was specifically observed in Ufm1- and Ufl1-deficient erythrocytes (Fig. 6C and S5D), while Mre11a deficiency resulted in telomere attrition in both erythroid and non-erythroid cells (Fig. 6D and S5E).

**Figure 6:**
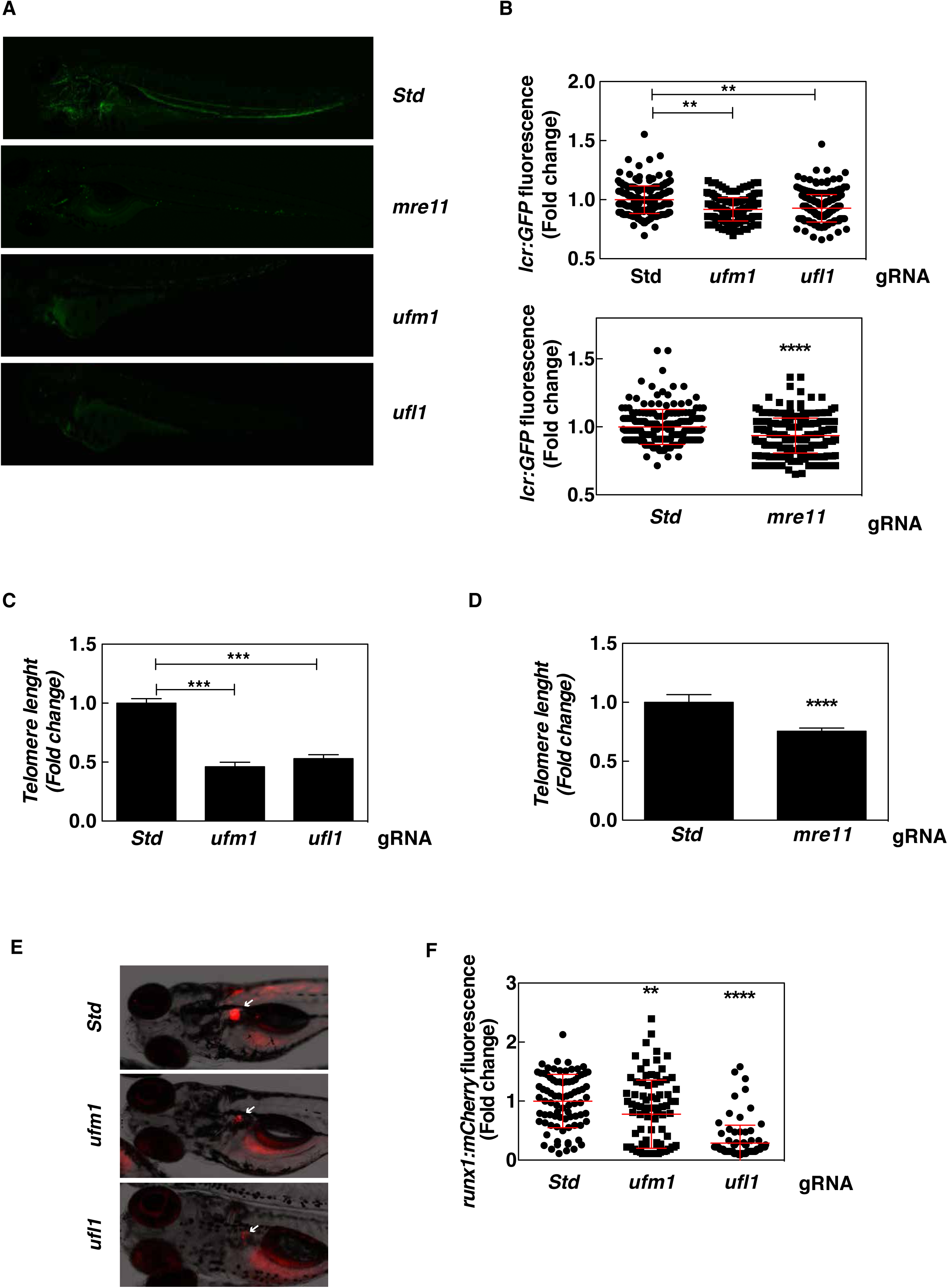
UFM1 pathway is requires for telomere stability and HSC survival *in vivo*. *Tg(lcr:eGFP)* (A-D) and *Tg(runx1:GAL4; UAS:nfsb-mCherry)* (E, F) one-cell embryos were injected with standard control (Std), *ufm1, ufl1* or *mre11a* sgRNA and recombinant Cas9. Representative images of green and red channel of whole larvae for the different treatments (A, E) and quantitation of erythroid cells at 4 dpf (B) and HSPCs at 5 dpf (F) are shown. Each dot represents normalized fluorescence from a single larva, while the mean ± SEM for each group is also shown. The sample size was: Std 228, *umf1* 182, *ufl1* 248 and Std 195, *mre11a* 251 (B); Std 83, *ufm1* 89, *ufl1* 91 (E). White arrows indicated HSPCs in the kidney marrow of 5 dpf larvae. (C, D) Telomere length was determined by qPCR in sorted erythroid cells (GFP^+^) of 6 dpf larvae. The data are shown as the mean±SEM of two independent experiments. **p<0.01; ***p<0.001; ****p<0.0001 according to Student *t* test (D) and ANOVA followed by Tukey multiple range test (B, C, F). See also Figure S5.

We then analyzed HSC emergence, maintenance, and differentiation using the lines *Tg(runx1:nfsB-mCherry)* that has fluorescently labeled hematopoietic stem and progenitor cells (HSPCs) (Tamplin *et al*, 2015), and *Tg(mpx:eGFP)* that marks neutrophils (Renshaw *et al*, 2006). Using these models, we found that HSPC survival was impaired in Ufm1- and Ufl1-deficient larvae after 3 dpf although the emergence of HSPCs (Fig. S5F) and primitive myelopoiesis (Fig. S5G) were both unaffected, Further, decreased number of HSPCs (Fig. 6E, F) and neutrophils (Fig. S5H) were observed at 5 dpf in the kidney marrow, the definitive hematopoietic tissues.

Taken together, these results convincingly show that disrupting the UFM1 pathway or MRE11 in a vertebrate animal model results in telomere attrition *in vivo*, with severe consequences for hematopoiesis.

## Discussion

Here we reveal a new and important role for the UFM1 pathway in telomere maintenance. We discovered the regulation of the TRF2-MRE11 interaction by the posttranslational modification of MRE11 with UFM1 and we further report that the severe defect in MRE11/TRF2 interaction observed in UFL1 KO HeLa cells is associated with marked telomeres shortening. We therefore link for the first time in mammals a telomere shortening phenotype with a lack of interaction between MRE11 and TRF2.

Intriguingly, the action of the MRN complex at telomeres have been essentially described when TRF2 is removed from the telomeres (Attwooll *et al*, 2009) (Deng *et al*, 2009) (Dimitrova & de Lange, 2009). This raises the question of the functional role of the established MRN-TRF2 interaction that occurs in S phase (Zhu *et al*, 2000) and that we find in this study regulated by UFM1 (this study). Interestingly, previous results reported that reduction of MRN by the use of siRNA, resulted in a transient shortening of G-overhang length in telomerase-positive cells suggesting that MRN promotes the formation of G-overhangs (Chai *et al*, 2006). The fact that reduction of overhang size observed in MRN deficient cells was observed upon expression of exogenous telomerase led to the proposal that the MRN complex might regulate telomerase action (Chai *et al*, 2006). While the mammalian MRN complex did not seem to have a marked effect on telomere length regulation (Attwooll *et al*, 2009), our work indicates that the lack of interaction between TRF2 and MRE11 caused by the absence of MRE11 UFMylation correlates with telomere shortening. One possibility consistent with Chai’s work would be that the interaction of UFM1-MRE11 to telomere via its interaction with TRF2 would favor the recruitment or action of telomerase through the regulation of G-overhangs. Interestingly, it was recently shown that phosphorylation of NBS1 triggered the dissociation of NBS1 from TRF2 (Rai *et al*, 2017). This dissociation in turn favors the interaction between TRF2 and Apollo/SNM1B (Rai *et al*, 2017) whose access to telomere regulates telomeric 3’ overhang (Wu *et al*, 2012). Whether MRE11 UFMylation impact Apollo/SNM1B access to telomere remains an open question. Another plausible hypothesis to explain the marked telomere shortening would be that MRE11 UFMylation promotes efficient replication of telomeric repeats to prevent fork collapse and brutal telomere shortening. Finally, we cannot exclude that *UFL1* knockout affects telomerase activity directly, either at the level of transcriptional downregulation of the *TERT* gene or telomerase complex assembly and intracellular trafficking.

The relevance of the signaling pathway uncovered in this study is demonstrated by the crucial role played by UFMylation in the maintenance of telomere length in HSCs in zebrafish. Our results suggest that UFMylation of Mre11 are dispensable for primitive myelopoiesis and emergence of HSPCs, but required for HSPC survival. In addition, the robust telomere attrition observed in erythrocytes of Ufl1- and Ufm1-deficient larvae, as well as in both erythroid and non-erythroid cells of Mre11a-deficient larvae, indicates that UFMylation of Mre11 is required to maintain telomere length and survival of HSPCs. Surprisingly, genetic inactivation of the main telomerase components, *tert* and *terc*, resulted in only modest telomere shortening in zebrafish larvae (Alcaraz-Pérez *et al*, 2014) compared with the inactivation of Ufl1, Ufm1 and Mre11a. This further bolsters our findings for the critical role played by UFMylation and Mre11 in telomere maintenance in HSC *in vivo*.

Some human tumors maintain their telomeres by a telomerase independent mechanism termed Alternative Lengthening of Telomeres (ALT). ALT involves Break Induced Replication (BIR) and requires the activity of the MRN complex (Zhong *et al*, 2007). We show here that UFL1 is not necessary for HR in telomerase-positive HeLa cells. This appears to be contradictory with the recent finding in ALT-dependent U2OS cells where UFL1 seems to be essential for HR and DSB repair (Qin *et al*, 2019). We explain this apparent discrepancy by hypothesizing that UFMylation of MRE11 could regulate HR specifically at telomeres. Therefore, cells relying on ALT for their telomere maintenance would have a more severe phenotype. This could explain why we did not manage to generate UFL1 KO in U2OS cells (data not shown). Additionally, it has been shown that UFMylation of MRE11 controls MRN complex formation (Wang *et al*, 2019). Since it has been shown that loss of the interaction between MRE11 and NBS1 is lethal for cells (Kim *et al*, 2017), the UFL1 KO cells should not be viable if the interaction between MRE11 and NBS1 is completely disrupted. This is consistent with what we show here, i.e, there is no impact on the MRN complex formation when UFL1 is deleted. In this context we can imagine that UFM1 regulates the formation of a fraction of the complex. This hypothesis is consistent with what has been shown, i.e. a reduction of the interaction and not a complete loss of the MRN complex formation (Wang *et al*, 2019). We could also elaborate on the possibility that MRE11 UFMylation takes place at specific locations (e.g. at telomeres), where it may affect MRN complex formation, which would account for only a fraction of total cellular MRN.

Furthermore, it is tempting to speculate that UFL1-dependent MRE11 UFMylation attenuates ATM activation (consistent with our data, Fig 4D) at telomeres undergoing replication thus restraining DDR activation due to transiently unprotected telomeres. We anticipate that the H4 mono-UFMylation-dependent mode of ATM activation at DSBs as proposed by Qin and colleagues (Qin *et al*, 2019) would not operate at telomeres for two reasons. Firstly, this mode of ATM activation relies uniquely on the recruitment of SUV39H1 to UFM1-H4, which then trimethylates H3K9. Importantly, SUV39H1 is absent from telomeres and H3K9me3 is deposited instead by SETDB1 (Gauchier *et al*, 2019). Secondly, ATM activation at telomeres is suppressed by telomere-bound protein TRF2 (Denchi & de Lange, 2007) (Okamoto *et al*, 2013). In this sense, our findings that MRE11 interaction with TRF2 is completely dependent on MRE11 UFMylation at K282/K283 and that ATM is constitutively active albeit at a low level in UFL1 KO HeLa cells is particularly intriguing. Since ATM promotes HR at DSBs (Beucher *et al*, 2009) (Bakr *et al*, 2015) and particularly at single-ended DSBs arising at broken replication forks (Balmus *et al*, 2019), attenuation of ATM activity at telomeres may be one way to suppress recombination at the forks that frequently stall and may break at repetitive and G-rich telomeric sequences. It is plausible that this mechanism is deregulated in the cells that maintain their telomeres via ALT, and this is likely the root of the differences between our results and previous studies.

In summary, here we identified a new mechanism regulated by an emerging posttranslational modification that is critical for, telomere length maintenance and HSC survival *in vivo*. We have identified the first MRE11 mutation impacts its interaction with TRF2. Studying the impact of this mutation on MRE11 function at telomere is likely to reveal fresh insights into the role of the MRN complex in the maintenance of telomere in mammals.

## Acknowledgements

This work was supported by the ATIP AVENIR, plan cancer (C17004AS), the canceropole PACA, the Spanish Ministry of Science, Universities and Innovation (grant BIO2017-84702-R to VM and PI13/0234 to MLC, co-funded with FEDER), and the University of Murcia (PhD fellowship to EMS). Work in VG’s Laboratory is supported by the “Ligue Nationale Contre le Cancer”, Equipe labellisée. We acknowledge Jean-Hugues Guervilly and Mauro Modesti for discussion, advices and comment, the two students in the lab who helped with this work, Lea Verrier and Leyla Ameur, the CRCM facilities including TrGET, microcopy and FACS, and I Fuentes and PJ Martínez for excellent technical assistance and zebrafish maintenance. Proteomics analyses were supported by the Institut Paoli-Calmettes and the Centre de Recherche en Cancérologie de Marseille. Proteomic analyses were done using the mass spectrometry facility of Marseille Proteomics (marseille-proteomique.univ-amu.fr) supported by IBISA (Infrastructures Biologie Santé et Agronomie), the Cancéropôle PACA, the Provence-Alpes-Côte d’Azur Région, the Institut Paoli-Calmettes, and the Fonds Européen de Développement Régional (FEDER).

## Authorship

LL and ABP designed and conducted most of the experiments. JP made in vitro ufmylation assay. EM made zebra fish experiment. GA quantified cell sensitivity. DR analyzed the interaction with TRF2. DC made TESLA and telomere Southern blot analyses. SA and LC made mass spectrometry experiments and data analysis. CL designed and supervised the study with VG, YK, VM and MLC. CL wrote the manuscript with input from YK, VM and VG.

## Materiel and methods

### Cell culture and transfection

293T and Hela FRT-T-Rex (Invitrogen) were maintained at 37 °C in DMEM (Dulbecco’s modified Eagle’s medium; Gibco) supplemented with 10% FBS, 1% penicillin/streptomycin (GIBCO). For transfection, each dish of adherent cells was transfected with 5–10 µg of plasmid DNA using lipofectamine 2000 (ThermoFisher).

### CRISPR gene editing

CRISPR Cas9 px335 and pBabe-U6 have been used to clone gRNA sense and antisense respectively. Cloning have been made according to Zhang lab protocols. Cells have been transfected with 1µg of combined plasmids and cell sorted into 96 well plates. Clones have been screened using immunodetection of targeted proteins and the mutation have been sequenced as described before (Munoz *et al*, 2014).

### Stable cell line establishment

Stably cells expressing BirA-GFP or BirA-tagged proteins were generated according to the manufacturer’s instructions. Briefly, pcDNA5 FRT BirA plasmids were co-transfected with POG44 plasmids (ratio 1/9) with lipofectamine 2000 (Invitrogen). Cell were then selected with Hygromycin and blasticidin.

### Cloning

Restriction enzyme digestions, DNA ligations and other recombinant DNA procedures were performed using standard protocols. All mutagenesis was performed using the QuikChange site-directed mutagenesis method (Stratagene) with KOD polymerase (Novagen). All DNA constructs were verified by DNA sequencing, which was performed by GATC (Eurofins). DNA for mammalian cell transfection was amplified in *E. coli* DH5α strain, and plasmid preparation was done using Qiagen Maxi prep Kit according to manufacturer’s protocol.

### Biotin ligase assay

Expression of BirA-proteins was induced for 24h by adding 1µM Doxycycline (sigma). For purification of biotinylated substrates, 10 × 10 cm dishes were treated for 8 hours with 50 µM biotin. Cells were then collected, washed with phosphate buffered saline (PBS) and resuspended in lysis buffer [0.5 ml/10 cm dish; 8 M urea, 1% SDS, 50 mM N-ethylmaleimide, 1× protease inhibitor cocktail (Roche) in PBS]. Sonication was performed as needed to reduce sample viscosity. To reduce urea concentration, the samples were diluted by adding binding buffer (3 M urea, 1 M NaCl, 0.25% SDS; 0.5 volume). Incubation was done using 100 µl suspension of high-capacity Streptavidine-sepharose beads (GE Healthcare) overnight at room temperature (RT). Washes have been done with 2× WB1, 3× WB2, 1× WB3, 3× WB4, 1× WB1, 1× WB5 and 3× WB6 [WB1: 8 M urea, 0.25% SDS in PBS; WB2: 6 M guanidine hydrochloride in PBS; WB3: 6.4 M urea, 1 M NaCl, 0.2% SDS in PBS (pre-warmed to 37 °C); WB4: 4 M urea, 1 M NaCl, 10% isopropanol, 10% ethanol, 0.2% SDS in PBS; WB5: 8 M urea, 1% SDS in PBS; WB6: 2% SDS in PBS]. Samples were eluted in 100 µl of 4× Laemmli sample buffer with 100 mM DTT by two cycles of heating (5 minutes; 99 °C), with vortexing in between. For MS analysis, the bead slurry was transferred to a Vivaclear Mini 0.8 µm PES filter (Sartorius) and spun to recover bead-free eluate.

### Mass spectrometry

Pulldown elutions were separated by SDS-PAGE and stained with Brilliant Blue G-Colloidal Concentrate (Sigma) according to manufacturer’s instructions. Gel bands were excised from the whole gel lane, destained and proteins were in-gel digested with trypsin (sequencing grade, Promega) overnight. The resulting peptide mixtures were analyzed by liquid chromatography (LC)-tandem mass spectrometry (MS/MS) using Orbitrap Fusion Lumos Tribrid Mass Spectrometer (Thermo Electron, Bremen, Germany) online with a nanoLC Ultimate 3000 chromatography system (Dionex, Sunnyvale, CA) through a LC EASY-Spray C18 column from Dionex. All raw LC-MS files were processed with MaxQuant software (version 1.5.3.8, www.maxquant.org) and search against species-specific Uniprot protein sequence databases and common contaminants using the Andromeda peptide search engine with a false discovery rate of 0.01 at both peptide and protein level

The lists of proteins identified by MS were analyzed as follows. First, contaminants and proteins identified by only one peptide were eliminated. Then, only those proteins with at least 10-fold higher peak surface in the experiment samples *versus* the controls BirA-GFP were considered as positive hits.

### Immunobloting

For immunobloting, 30µg of total protein extracted with lysis Buffer (50 mM Tris–HCl (pH 7.5), 150 mM NaCl, 1 mM EDTA, 1 mM EGTA, 1% (w/v) Triton, 1 mM sodium orthovanadate, 10 mM sodium glycerophosphate, 50 mM sodium fluoride, 10 mM sodium pyrophosphate, 0.27 M sucrose, 0.1% (v/v) 2-mercaptoethanol, 1 mM benzamidine and 0.1 mM PMSF) or 25µl of protein fraction (Subcellular protein fractionation kit, ThermoFisher) have been separated by SDS PAGE 4-12% (Bis-tris Bolt, ThermoFisher), transferred on nitrocellulose (Turbo transfer Bio-Rad). The signal was detected with chemidoc (Bio-Rad). Larvae of 6dpf were lysed in lysis buffer (50 mM Tris-HCl, pH 7,5, 1% IGEPAL, 150 mM NaCl and a 1:20 dilution of the protease inhibitor cocktail P8340 from Sigma Aldrich). Samples were centrifuged (13.000 × g, 10 min) and resolved on 4-12% SDS-PAGE and transferred to PVDF membranes. Membranes were block 1h at RT and incubated overnight at 4°C. For detection, corresponding horseradish peroxidase conjugated secondary antibodies (1:5000 dilution) were used. After repeated washes, the signal was detected with the enhanced chemiluminescence reagent and ChemiDoc XRS Bio-Rad.

### Immunoprecipitation

For Immunoprecipitation, of GFP, TRAP-GFP beads were used. 0.5–5 mg of lysates was incubated with 10–20 µl of resin for 2 h at 4°C under gentle agitation, and the immunoprecipitates were washed three times with lysis buffer containing 0.15 M NaCl and then twice with lysis buffer. Proteins were eluted by resuspending washed immunoprecipitates in 30 µl of 1× SDS sample buffer.

### Immunofluorescence

Cells were washed with PBS and fixed for 20 min with 2% PFA in PBS. Cells were permeabilised for 5 min with PBS/0.2% Triton X-100, and blocked with PBS/0.2% Tween 20 (PBS-T) containing 5% BSA. Coverslips were incubated for 1 h with primary antibodies and for 30 min with appropriate secondary antibodies coupled to Alexa Fluor 488 or 594 fluorophores (Life Technologies), before being incubated with 2µg/ml DAPI. Pictures were acquired with Z1 (Zeiss). For high resolution imaging, z-stacks were acquired with a Z1 (z-stack of 0.2 µm interval) equipped with a 63x oil objective (ZEISS) and controlled with Zen. Deconvolutions were then performed in conservative mode. The different channels were acquired sequentially.

### Bacterial protein purification

Recombinant His_6_-Uba5, His_6_-Ufm1, His_6_-MRE11 Wild type and mutants were expressed in *E.coli* BL21 and purified using Ni^2+^-NTA affinity chromatography. Briefly, *E.coli* BL21 cultures expressing His_6_-tagged proteins were grown in 2xTY medium at 37°C until OD 0.6. Protein expression was induced with 0.5 mM IPTG and the cultures were incubated at 18 C for 16 hrs. Cells were lysed in lysis buffer (25 mM Tris 7.5, 300 mM NaCl, 10% Glycerol, 2 mM DTT,1 mM Benzamidine, 1 mM AEBSF, protease inhibitor cocktails (Roche)). Lysed cells were then clarified by centrifugation at 30000xg for 30 mins at 4°C. The clarified lysate was then incubated with Ni^2+^-NTA Agarose beads for 2 hrs in binding buffer (25mM Tris 7.5, 300 mM NaCl, 10% Glycerol, 10 mM Immidazole), washed extensively and eluted using binding buffer with 300 mM Immidazole. His_6_-tags were cleaved off by incubating tagged proteins with C3 protease at 4°C overnight. GST-tagged Ufc1 was expressed in *E.coli* BL21 as described above. Cells were lysed in lysis buffer containing 25mM Tris 7.5, 300mM NaCl,10% Glycerol and 2mM DTT using ultrasonication. Glutathione B Sepharose beads were incubated with clarified lysate for 2 hrs. The beads were then washed with high salt buffer containing 25 mM Tris-HCl pH 7.5, 500 mM NaCl, 10% glycerol and 1 mM DTT. The beads were washed with low salt buffer containing 25 mM Tris-HCl pH 7.5, 500 mM NaCl, 10% glycerol and 1 mM DTT. The GST tag was then cleaved off by incubation with C3 protease at 4°C overnight. All proteins were further purified by size exclusion chromatography using Superdex 75 16/60 and Superdex 200 16/60 columns (GE Healthcare Life Sciences). The purified proteins were then concentrated and stored at −80°C.

### In vitro Ufmylation assay

Recombinant fragments of MRE11 and mutants were incubated with 0.25µM Uba5, 5µM Ufc1, 2µM Ufl1 and 30µM Ufm1 in reaction buffer containing 50 mM HEPES 7.5, 10 mM MgCl_2_ and 5 mM ATP for 1 hr at 37°C. The reaction was stopped by the addition of 3x SDS loading buffer containing 10% mercaptoethanol. The reaction products were separated on a 4-12% NuPAGE SDS-PAGE gel and analysed by Immunoblotting using specific MRE11 antibodies.

### Cell sensitivity assay

For sensitivity assay to IR, 1000 cells were plated in three replicates onto 10 cm plates in complete growth medium. After cells attached, they were treated with indicated dose of IR. The number of colonies with > 100 cells was counted. For each genotype, cell viability of untreated cells was defined as 100%. Data are represented as mean ± SD from three independent experiments. For the other cytotoxicity assays, 1000 cells were plated 96 well dishes. Next day, indicated drugs were added to the wells and plates were transferred into an IncuCyte microscope (Essen BioScience). Phase contrast pictures were acquired every 3 h over 48 h. Percentage of cell confluence was calculated by the Cell Player integrated software (Essen BioScience) and analysed with GraphPad Prism® software.

### Homologous recombination assay

Hela and Hela UFL1 KO were transfected with 5 µg of pCBA-I-SceI and 5 µg of DR-GFP. 24h later, cells were harvested and analyzed by fluorescence-activated flow cytometry (FACS) to examine GFP positive cells. Cells were gated to exclude cellular aggregates, debris and GFP negative cells in the FSC/FSC dot plot. Gates of positive cells were set and compared with a control sample (without pCBA-I-SCEI). Results were normalized with transfection efficiency using mCHerry plasmids.

### Zebrafish experiments

For crispR experiments zebrafish lines used were *Tg(mpx:eGFP)*^*i114*^ (Renshaw et al., 2006), *Tg(lcr:eGFP)*^*cz3325*^ (Ganis et al., 2012), *Tg(runx1:GAL4)*^*utn6*^ (Tamplin et al., 2015) and *Tg(UAS:nfsB-mCherry)*^*c264*^ (PMID: 17335798). The sgRNAs were obtained from IDT® and prepared used the manufacturer’s manual with a concentration of resulting duplex of ∼1715ng/uL (50uM). After Assembling the Ribonucleoprotein Complex 1 nL of the mix was injected into the yolk of 1-cell stage zebrafish embryos. The sequences of the guides are: *ufm1* 5′-TGAGAGCACACCATTCACAG CGG-3′, *ufl1* 5′-CCC AGAGCACTTGGGTTGAGTCG-3′, *mre11a* 5′-GGCAACCATGATGACCCAAC TGG-3′.

Images were acquired at 3, 4 and 5 dpf using a Leica M205 FA fluorescence stereo microscope equipped with a DFC365FX camera (Leica) and processed using ImageJ software (http://rsb.info.nih.gov/ij/) and Photoshop CS.

Approximately 300 to 500 larvae were anesthetized in tricaine, minced with a razor blade, incubated at 28°C for 30 min with 0.077 mg/ml Liberase (Roche) and the resulting cell suspension passed through a 40 µm cell strainer. Cell sorting was performed on a SH800Z Cell Sorter (Sony).

### Telomere measurement by qPCR

Genomic DNA (gDNA) was extracted from the cells using the “ChargeSwitch™ gDNA Micro Tissue Kit (Invitrogen). The telomeric sequences were detected through real-time PCR using 16ng of gDNA as a template. Ribosomal protein S11 (rps11) content in each sample was used for normalization of zebrafish mRNA expression, using the comparative Ct method (2-DCt). Actin and 36b4 were used as a standard value for lymphocytes and HeLa cell line, respectively. Reaction mixtures were incubated for 15 min at 95 °C, followed by 40 cycles of 15 s at 95 °C, 2 min at 54 °C, and finally 15 s at 95 °C, 1 min at 60 °C, and 15 s at 95 °C. For the standard genes reaction mixtures were incubated for 10 min at 95 °C, followed by 40 cycles of 15 s at 95 °C, 1 min at 60 °C, and finally 15 s at 95 °C, 1 min at 60 °C, and 15 s at 95 °C.

The primers used were: Telom F: 5’-TTTTTGAGGGTGAGGGTGAGGGTGAGGGTGAGGGT-3’ and Telom R: 5’-TCCCGACTATCCCTATCCCTATCCCTATCCCTATCCCTA-3’. rps11 F:5’-ACAGAAATGCCCCTTCACTG-3’ and rps11 R: 5’-GCCTCTTCTCAAAACGGTTG-3’. 36b4 F: 5’-CAGCAAGTGGGAAGGTGTAATCC-3’ and 36b4 R: 5’-CCCATTCTATCATCAACGGGTACAA-3’. Actin F: 5’-GGCACCACACCTTCTACAATG-3’ and R: 5’-GTGGTGGTGAAGCTGTAGCC-3’.

The results were normalized with the control. In all cases, each PCR was performed with triplicate samples and repeated at least with two independent samples. The differences between two samples were analysed by the Student’s t-test and between three samples by One-way ANOVA.

### Metaphase spread analysis

Cells were initially plated in DMEM containing 10% FBS until cells reached 60–70% confluence. All cultures were harvested following conventional cytogenetic protocol (Brown & Lawce, 1997). Briefly, the cell cultures were treated with 0.1 µg/ml Colcemid (Irvine Scientific) approximately 30 min before the initiation of harvest. For chromosome preparations, the cells were harvested following conventional cytogenetic protocol of hypotonic treatment (75 mM KCl) and freshly prepared chilled 3:1 methanol:acetic acid fixation. This was followed by four additional fixation cycles and air-dried slide preparation. The slides were ‘aged’ in a hot oven at 60°C over 16 h, followed by Giemsa staining (Invitrogen). A total of 25 metaphases were scored for each culture.

### Analysis of cell cycle by flow cytometry

Cells were analyzed for their respective cell cycle phase distribution using flow cytometry. Cells were trypsinised, washed with PBS + 0.2% (w/v) BSA and resuspended in flow cytometry tubes. Cells were then fixed by 70% (v/v) ice-cold ethanol and stored at −20°C until analysis. After washing fixed cells once with PBS, RNase A (50 µg/ml) and PI (50 µg/ml) were added to the cells and incubated in the dark at room temperature (25°C) for 20 min. The live cell populations were then subjected to quantitative measurement of DNA content by flow cytometry using a LSRFortessa (BD Biosciences) and cell cycle distribution and the percentage of G2–S–G1 cells determined by the Watson (pragmatic) modelling algorithm using FlowJo software (Treestar).

### Telomere FISH

Telomere FISH on metaphase chromosome spreads was performed according to standard protocol (Lansdorp et al., *Hum Mol Gen* 1996). Briefly, a hybridization mix containing 50 nM Cy3-labeled PNA telomere probe (Cy3-OO-TTAGGGTTAGGGTTAGGG 3’) in the 70% formamide was spotted on the slides with metaphase spreads prepared as described above. The slides were covered with coverslips, pre-heated to 80°C to facilitate DNA denaturation, and were incubated at room temperature in the dark for a minimum of 2-3 hours to allow PNA probe annealing to telomeres. Following hybridization, the slides were washed two times, 15 min each, with the solution containing 10 mM Tris-HCl pH 7.2, 70% formamide, and 0.1% BSA; and then three times, 5 min each, with the solution containing 0.1 M Tris-HCl pH 7.2, 0.15 M NaCl; 0.08% Tween-20. After washes, the chromosome spreads were dehydrated in ethanol series: 5 min in 70%, 95%, 100% ethanol, air-dried, and sealed using a coverslip and a small volume of an antifade solution containing DAPI (4′, 6-diamidino-2-phenylindole) as a DNA counterstain. The images were acquired using Zeiss fluorescence microscope and the Metamorph software as described above. The PNA probe was ordered from PNA Bio Inc (Thousand Oaks, CA, USA), dissolved at 50 µM in formamide, and stored at −80°C.

### Southern blot analysis of terminal restriction fragments (TRFs)

The cells were lysed in the TNES buffer (10 mM Tris-HCl pH 7.5, 400 mM NaCl, 100 mM EDTA, 0.6 % SDS), supplemented with Proteinase K (100 µg/ml), and incubated at 55°C for 2 hours. Genomic DNA was extracted using phenol:chloroform (1:1) mixture, precipitated with 100% ethanol-acetate, washed with 70% ethanol, dissolved in the 10 mM Tris-HCl pH 8.0, and quantified using NanoDrop spectrophotometer. The purified DNA (12 µg of each sample) was digested with *Hin*fI and *Rsa* I (1.5 U each per µg of DNA) at 37°C overnight and separated on the 0.8% agarose gel in Tris-borate buffer. The DNA was transferred from the gel on the Hybond N+ membrane using the standard Southern blotting procedure in the alkaline solution (0.4 N NaOH; 0.5M NaCl). The DNA was UV-crosslinked onto the nylon membrane and then blocked and hybridized with ^32^P-labeled (CCCTAA)_3_ probe in the Church and Gilbert buffer overnight at 42°C. After washing in the Na-phosphate buffer, the membrane was exposed to phosphoimager screen for image acquisition.

### Telomere Shortest Length Assay (TeSLA)

TeSLA was performed according to the protocol described by Lai et al. (Nat Comm 2017). Briefly, 50 ng of undigested genomic DNA was ligated with an equimolar mixture (50 pM each) of the six TeSLA-T oligonucleotides containing seven nucleotides of telomeric C-rich repeats at the 3′ end and 22 nucleotides of the unique sequence at the 5’ end. After overnight ligation at 35°C, genomic DNA was digested with *Cvi*AII, *Bfa*I, *Nde*I, and *Mse*I, the restriction enzymes that create short either AT or TA overhangs. Digested DNA was then treated with Shrimp Alkaline Phosphatase to remove 5′ phosphate from each DNA fragment to avoid their ligation to each other during the subsequent step of adapter ligation. Upon heat-inactivation of phosphatase, partially double-stranded AT and TA adapters were added (final concentration 1 µM each) and ligated to the dephosphorylated fragments of genomic DNA at 16°C overnight. Following ligation of the adapters, genomic DNA was diluted to a final concentration 20 pg/µL, and 2-4 µL of it was used in a 25 µL PCR reaction to amplify terminal fragments using primers complementary to the unique sequences at the 5’ ends of the TeSLA-T oligonucleotides and the AT/TA adapters. FailSafe polymerase mix (Epicenter) with 1× FailSafe buffer H was used to efficiently amplify G-rich telomeric sequences. Entire PCR reactions were then loaded onto the 0.85% agarose gel for separation of the amplified fragments. To specifically visualize telomeric fragments, the DNA was transferred from the gel onto the nylon membrane by Southern blotting procedure and hybridized with the ^32^P-labeled (CCCTAA)_3_ probe essentially as described above for the Southern blot analysis of TRFs. The sizes of the telomeric fragments were quantified using TeSLA Quant software (Lai et al., Nat Comm 2017).

## Figure legends

**Figure S1:**
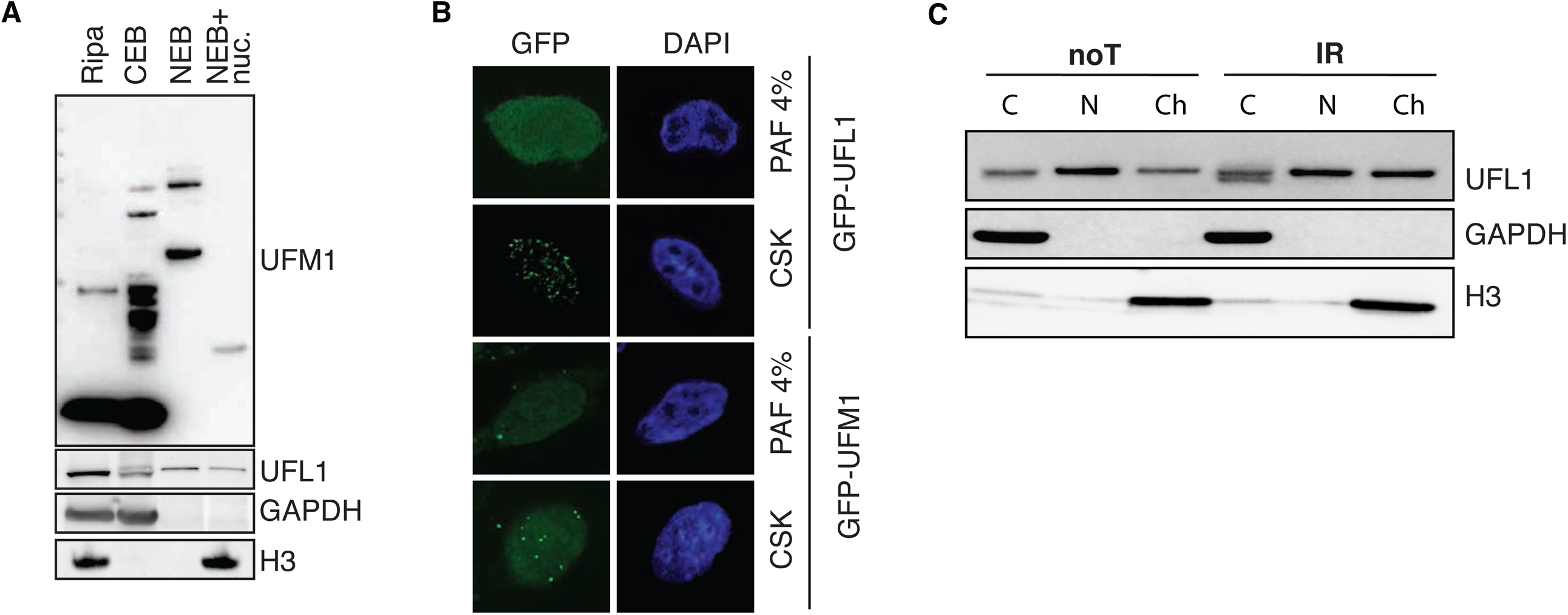
UFL1 localizes at the chromatin. (A) Immunoblot of UFM1, UFL1, H3 and GAPDH. C = cytoplasm fraction, N = Nuclear fraction and Ch= chromatin fraction. (B) GFP immunostaining of cells transfected with GFPUFL1 or GFP-UFM1. Cells were fixed with paraformaldehyde (PAF) or treated with CSK prior fixation to release soluble proteins. (C) Immunoblot of UFL1, H3 and GAPDH in the absence or presence of IR. C = cytoplasm fraction, N = Nuclear fraction and Ch= chromatin fraction.

**Figure S2:**
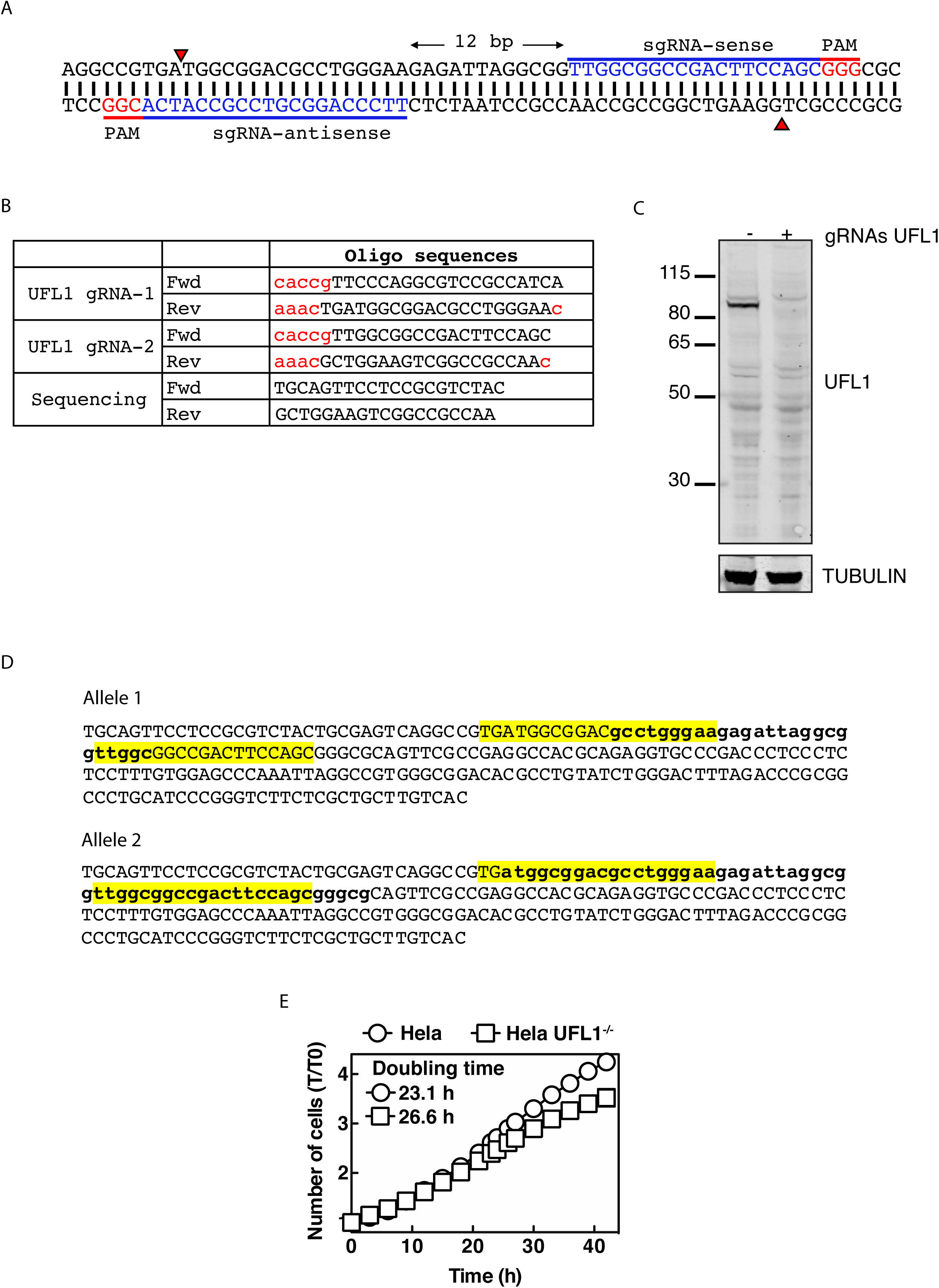
Generation of UFL1 KO in Hela cells. (A) Targeted site. (B) Sequences of Oligo used to generate and sequence the KO. (C) UFL1 immunoblot in WT and UFL1 KO cells. (D) DNA sequences of targeted region. (E) IncuCyte cell proliferation assay showing the growth WT and UFL1 KO cells up to 46 h.

**Figure S3:**
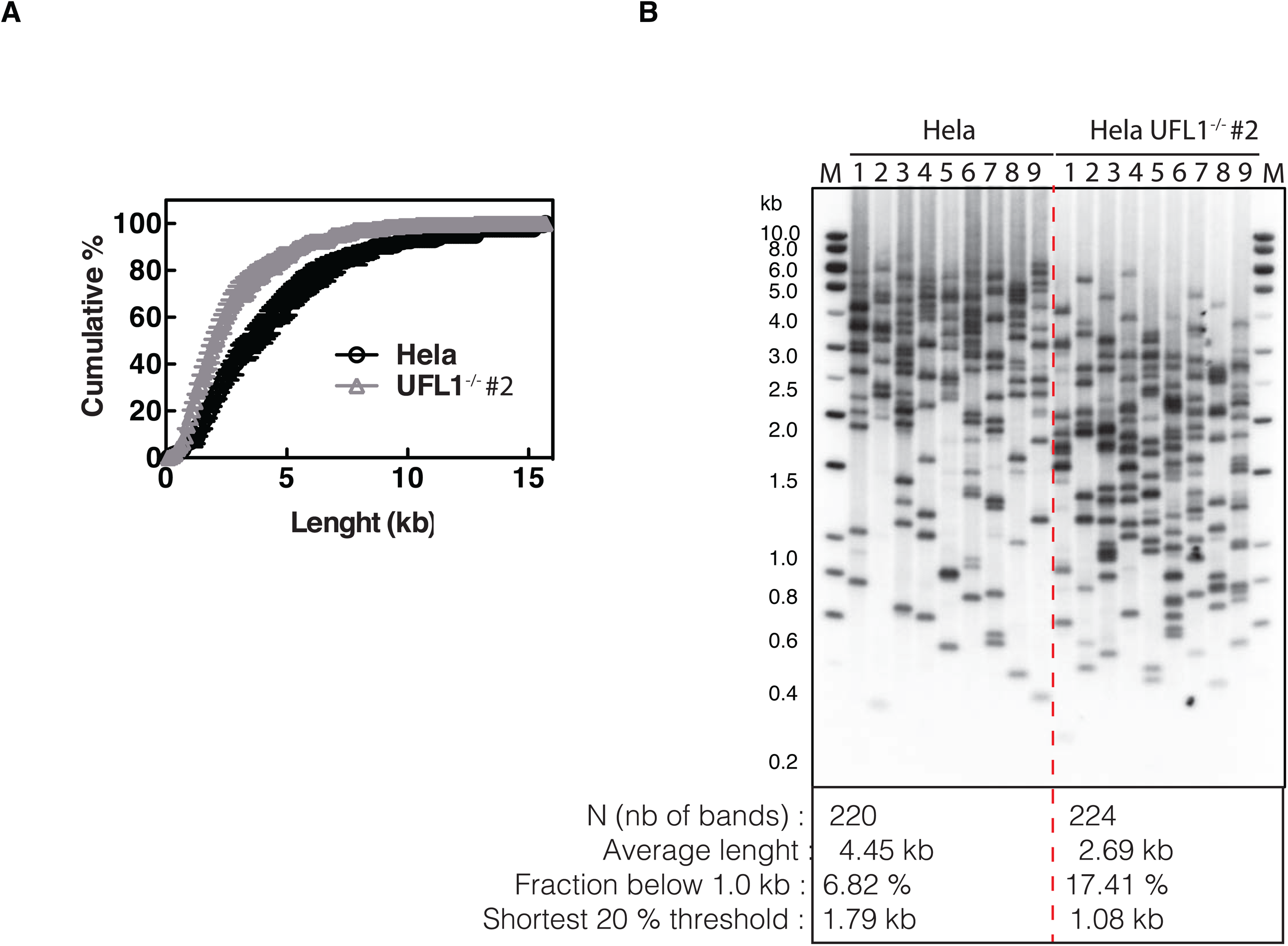
UFL1 KO cells have short telomeres. (A) Quantification of B. (B) Results of TeSLA using DNA as indicated. Nine TeSLA PCRs were performed for each DNA sample.

**Figure S4:**
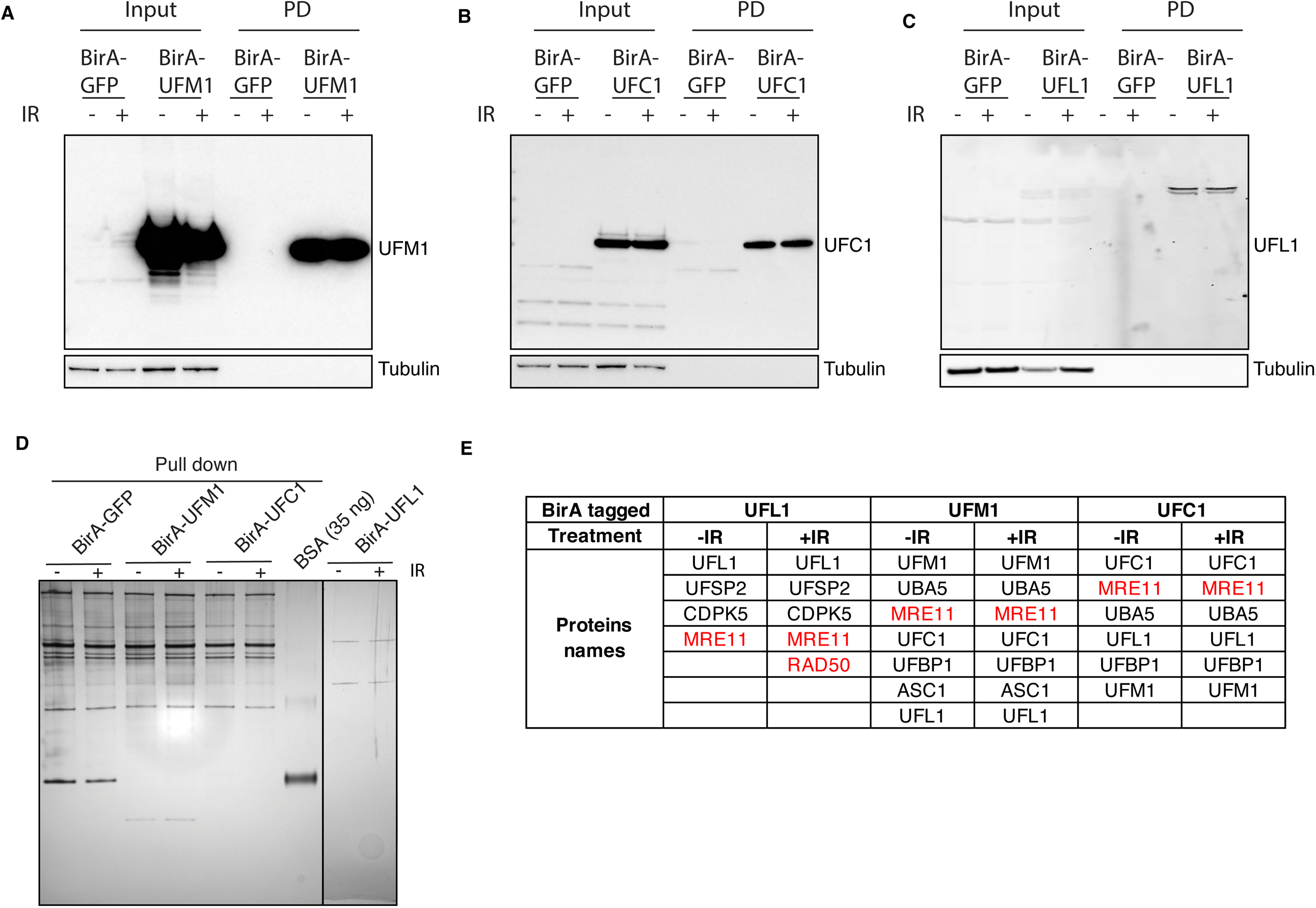
MRE11 interacts with the UFM1 pathway. (A) Biotinylated proteins are purified from cells expressing BirA-GFP or BirA-UFM1 with or without treatment with IR. The samples were then subjected to immunoblot with indicated antibodies. (B) Biotinylated proteins are purified from cells expressing BirA-GFP or BirA-UFC1 with or without treatment with IR. The samples were then subjected to immunoblot with indicated antibodies (C) Biotinylated proteins are purified from cells expressing BirA-GFP or BirA-UFL1 with or without treatment with IR. The samples were then subjected to immunoblot with indicated antibodies. (D) silver straining of the proteins purified in A-C. (E) List of the candidate target proteins identified by mass spectrometry.

**Figure S5:**
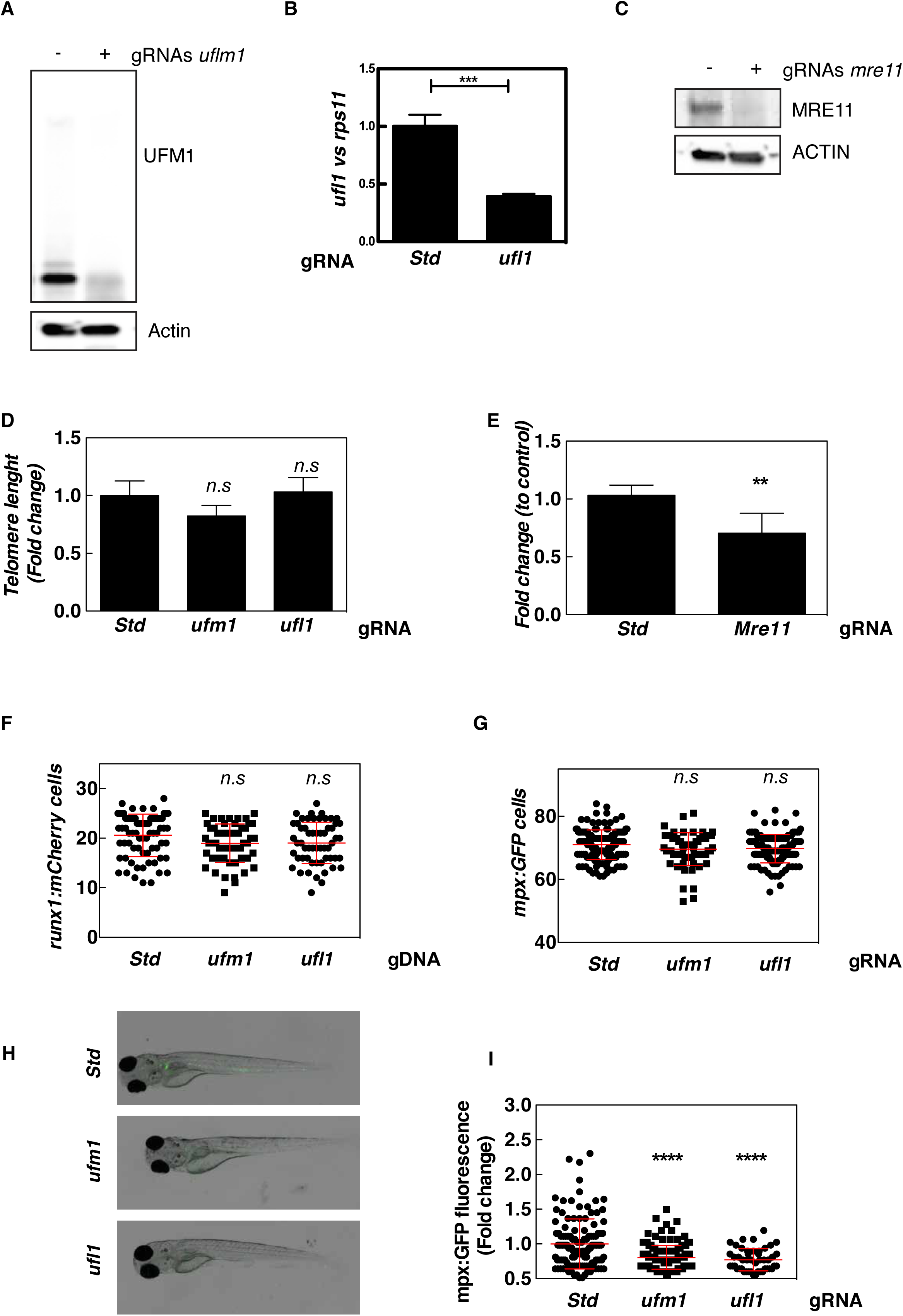
(Related to Figure 6): UFM1 pathway is dispensable for primitive myelopoiesis in zebrafish. *Tg(lcr:eGFP)* (A-E) and *Tg(mpx1:eGFP)* (H, I) one-cell embryos were injected with standard control (Std), *ufm1, ufl1* or *mre11a* sgRNA and recombinant Cas9. (A-C) The efficacy of the sgRNA was determined by western blot (A, C) or RT-qPCR (B) using Actb and *rps11* expression for normalization, respectively. One representative experiment is shown. (D, E) Telomere length was determined by qPCR in sorted cells (GFP^−^) of 6 dpf larvae. The data are shown as the mean±SEM of two independent experiments. (F-I) Quantitation of HSPCs (F) and neutrophils (G) at 2 dpf are shown. Each dot represents normalized fluorescence from a single larva, while the mean ± SEM for each group is also shown. The sample size was: Std 74, *ufm1* 55, *ufl1* 57 (F); Std 149, *ufm1* 70, *ufl1* 129 (G); Std 179, *ufm1* 174, *ufl1* 97. (I). Representative images of green channel of whole *Tg(mpx1:eGFP)* larvae for the different treatments are also shown (H). White arrows indicated neutrophils in the kidney marrow of 5 dpf larvae. n.s, non-significant; **p<0.01; ***p<0.001; ****p<0.0001 according to Student *t* test (B, E) and ANOVA followed by Tukey multiple range test (D, F, G, I). See also Figure S5.

